# Is resting state fMRI better than individual characteristics at predicting cognition?

**DOI:** 10.1101/2023.02.18.529076

**Authors:** Amir Omidvarnia, Leonard Sasse, Daouia I. Larabi, Federico Raimondo, Felix Hoffstaedter, Jan Kasper, Juergen Dukart, Marvin Petersen, Bastian Cheng, Götz Thomalla, Simon B. Eickhoff, Kaustubh R. Patil

## Abstract

Changes in spontaneous brain activity at rest provide rich information about behavior and cognition. The mathematical properties of resting-state functional magnetic resonance imaging (rsfMRI) are a depiction of brain function and are frequently used to predict cognitive phenotypes. Individual characteristics such as age, gender, and total intracranial volume (TIV) play an important role in predictive modeling of rsfMRI (for example, as “confounders” in many cases). It is unclear, however, to what extent rsfMRI carries independent information from the individual characteristics that is able to predict cognitive phenotypes. Here, we used predictive modeling to thoroughly examine the predictability of four cognitive phenotypes in 20,000 healthy UK Biobank subjects. We extracted common rsfMRI features of functional brain connectivity (FC) and temporal complexity (TC). We assessed the ability of these features to predict outcomes in the presence and absence of age, gender, and TIV. Additionally, we assessed the predictiveness of age, gender, and TIV only. We find TC and FC features to perform comparably with regard to predicting cognitive phenotypes. As compared to rsfMRI features, individual characteristics provide systematically better predictions with smaller sample sizes and, to some extent, in larger cohorts. It is also consistent across different levels of inherent temporal noise in rsfMRI. Our results suggest that when the objective is to perform cognitive predictions as opposed to understanding the relationship between brain and behavior, individual characteristics are more applicable than rsfMRI features.

---

Resting-state functional magnetic resonance imaging (rsfMRI) is widely used for studying human brain function [1]–[3]. Functional connectivity (FC) is an important aspect of rsfMRI defined as the statistical dependence between different brain areas during periods of rest or low cognitive demand [4]. A common application of rsfMRI is the prediction of cognitive performance [5]–[9], and clinical phenotypes [5], [10]–[13] [14]–[20]. This is usually accomplished by extracting various features from rsfMRI, such as widely used FC measures, and using them for predictive modeling. This approach has been boosted by modern MRI scanners with powerful magnetic field strengths, large databases, high-performance computing systems, computational software packages, and improved machine learning techniques [21]. However, the field has struggled to advance to real-world applications due to systematic challenges such as modest prediction accuracy in large populations (*N_subject_* > 2000) [5], [6], [22], [23] and replication failures of studies with small sample sizes [23]. An effective improvement could be to consider features beyond prevalent FC measures.

In the current rsfMRI-based prediction pipelines, individual characteristics such as age, gender, and total intracranial volume (TIV) are typically treated as “confounds’’ and hence removed from the rsfMRI features or from the prediction targets [24]. The rationale behind this practice is that any information other than that directly related to brain activity should be discarded because it may prevent us from determining the neuronal origin of the predictive signal [25]. However, there is evidence that features based on individual characteristics might be better at predicting measures of mental health than those based on fMRI [26], [27]. It remains unclear whether rsfMRI is more accurate than individual characteristics at predicting cognitive phenotypes. The answer to this question will help the scientific community better understand the potential and limitations of rsfMRI in cognitive prediction, and guide future research efforts to improve the accuracy and reliability of this technique. To investigate this, we developed four cognitive prediction scenarios using a wide range of rsfMRI features in conjunction with three typically considered confounds, age, gender, and TIV. We used a large sample from the UK Biobank (*N_subject_* = 20,000) that included high-resolution rsfMRI and four cognitive phenotypes: fluid intelligence, processing speed, visual memory, and numerical memory [28]. The cognitive phenotypes were then predicted using different rsfMRI features at the brain region of interest (ROI) level.

Functional connectivity and large-scale nonlinear interactions between brain regions are intertwined [29], [30]. The nonlinearity of functional connections at micro-, meso-, and macro-scales gives rise to a temporally complex (TC) behavior in the hemodynamic response of the brain across time, as measured by fMRI [31], [32]. There is evidence that the TC of rsfMRI and cognitive phenotypes are correlated [33]–[36], providing a promising feature space for brain-behavior predictions [33], [34], [37]–[42]. As TC collects different rsfMRI characteristics than FC, it is expected to be able to supplement FC’s prediction capacity. It is also unclear how these rsfMRI-derived properties relate to the “noise” profile of various brain regions, including head movement, heartbeat, and respiration [43]. In the cognitive prediction pipelines of this study, we utilized nine rsfMRI features covering five prominent characteristics of FC (*fractional amplitude of low-frequency fluctuations* or *fALFF* [44]*, local correlation* or *LCOR* [45]*, global correlation* or *GCOR* [46]*, Eigenvector centrality* or *EC* [47], and *weighted clustering coefficient* or *wCC* [47]) as well as four TC metrics (*Hurst exponent* or *HE* [48]*, Weighted permutation entropy* or *wPE* [49], *Range entropy* or *RangeEn* [50], and *Multiscale entropy* or *MSE* [51]). We then entered these features into the predictive modeling pipelines, considering different roles for age, gender, and TIV as follows: (*i*) rsfMRI features without removing age, gender, and TIV, (*ii*) rsfMRI features after removing of these individual characteristics (i.e., treating them as confounds), (*iii*) a combination of rsfMRI features with age, gender, and TIV, and (*iv*) age, gender, and TIV only. We also investigated the impact of ROI-wise temporal signal to noise ratio (tSNR) on the models to examine the influence of “noisy” brain regions in the predictions.

We found that despite describing different aspects of rsfMRI, FC and TC features have comparable predictive capacity. When comparing some pairs of rsfMRI features from the same subject in a sample, there was a high probability of correctly identifying which feature belongs to which subject. This indicates that the rsfMRI features are capturing individual-level patterns that are distinct enough to allow for accurate identification, even though they may be conceptually different. The removal of age, gender, and TIV from the features or targets resulted in reduced performance. In most cases, the sole use of age, gender, and TIV yielded the best prediction accuracy. The accuracy of the predictions improved marginally when using a combination of rsfMRI features and individual characteristics. The results also show that age and gender could be predicted more accurately than cognitive phenotypes in general. Our findings demonstrate that rsfMRI features may not be necessarily better than age, gender, and TIV at predicting cognitive phenotypes, even when the sample size is increased to large numbers.

## Results

### Quantifying rsfMRI complex dynamics and cognitive phenotypes

We used preprocessed rsfMRI data from 20,000 unrelated UK Biobank participants [52] and extracted four TC measures as well as five FC-derived measures from them (see Methods). As prediction targets, we chose the four most reliable cognitive phenotypes in the UK Biobank database, fluid intelligence, processing speed, visual memory, and numerical memory [53]. See Table S1 in the Supplementary Materials for more details.

We used ridge regression with l2-norm regularization for predictive modeling, a widely used model for cognitive phenotypic prediction using rsfMRI [5], [54], [55]. Model performance was measured through cross-validation using the Spearman correlation between the real and predicted targets in regression tasks or the balanced accuracy in classification tasks. We used nested-cross validation where the hyper-parameter α was tuned in the inner loop [27]. The impact of age, gender, and TIV, was addressed through four scenarios, outlined in Figures 1 (see also Methods).

**Figure 1:**
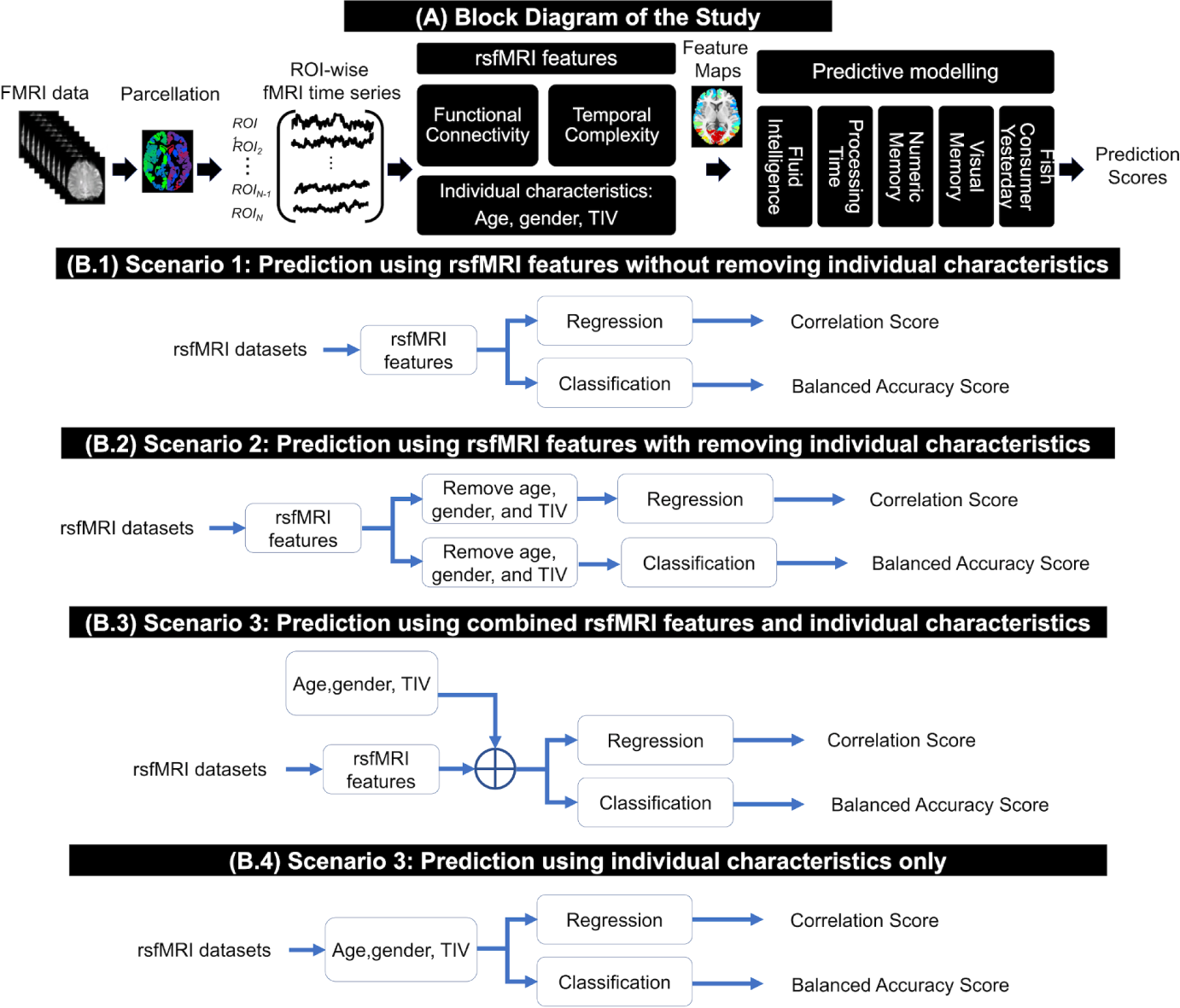
(A): main block diagram of this study, including the rsfMRI features and the prediction targets from the UK Biobank. (B) Four analysis scenarios based on the role of individual characteristics, i.e., age, gender, and total intracranial volume (TIV), in cognitive phenotypic prediction.

### Larger sample sizes increase accuracy but eventually reach a plateau

First, we examined whether increasing the sample size could improve the prediction accuracy of cognitive and non-cognitive targets in all four scenarios (Figure 1). Increasing the number of subjects improved accuracy most of the time, but the performance curves reached a plateau when using approximately more than 2,000 participants (Figure 2). As a sanity check, we tested all the predictive modeling scenarios using fish consumption (the day prior to fMRI) as a target presumably unrelated to the rsfMRI features. The performance for all sample sizes, rsfMRI features, individual characteristics, and their combinations remained at chance level (Figure 2).

**Figure 2:**
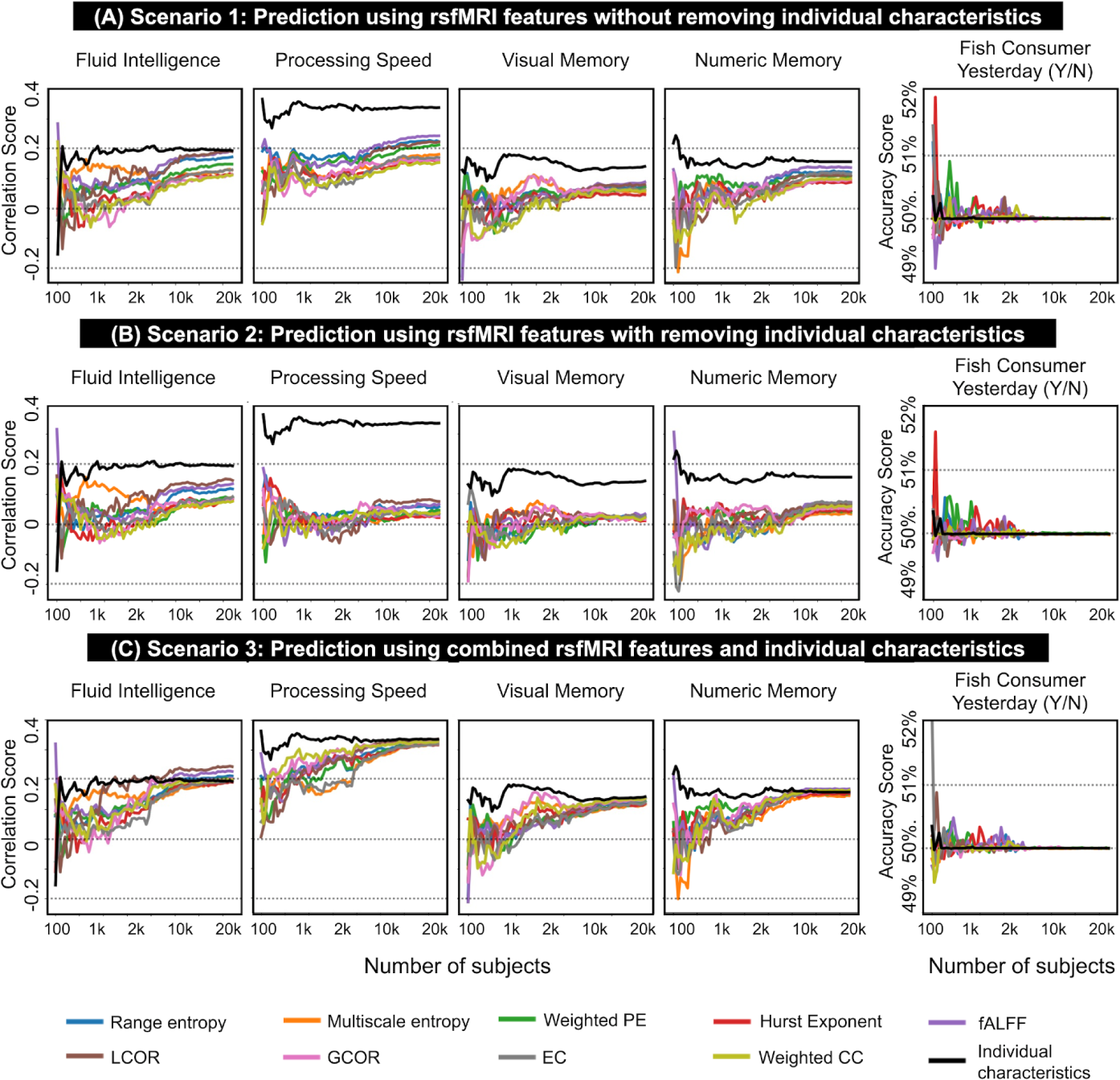
Prediction accuracy scores associated with nine rsfMRI features and five prediction targets using scenarios 1–3 of this study (see also Figures 1-B.1–B.3 and Methods). The prediction accuracies of individual characteristics only (Scenario 4 in Figure 1-B.4) have been plotted in bold black on all panels. Prediction accuracies of the fluid intelligence, processing speed, visual memory, and numeric memory scores are computed as the Pearson correlation between the actual values and predicted values through ridge regression modeling. The prediction accuracy of Fish consumer yesterday is computed as the balanced accuracy through ridge binary classification. Each rsfMRI feature is illustrated in a distinct color and listed in the figure legend. In each figure panel, the x-axis represents the population size in the analysis, and the y-axis shows the prediction accuracy. The predictive modeling of each pair of features and targets is repeated for different sample sizes in the UK Biobank, ranging from *N_subject_* = 100 to *N_subject_* = 20,000. The population sizes from 100 to 2000 were increased with a 50-step increment (see the light orange shadow in the figure panels) and from 2000 to 20,000 with a 500-step increment (see the light blue shadow in the figure panels). As the panels show, the prediction accuracy improved with increasing sample size and the number of suprathreshold ROIs (i.e., higher accuracy toward the upper right corner of the color-coded maps). See Supplementary Figure S1 for the boxplot representation of these results.

### TC and FC features show comparable predictive capacities

Next, we investigated how TC and FC features compare in cognitive phenotype prediction across different sample sizes. The average performance of ridge regression models across 5 repeats and 5 folds of nested cross-validation suggested that certain features, specifically *fALFF*, *LCOR*, *wPE*, and *RangeEn_B_*, performed better than others in all contexts, regardless of the target. The correlation between actual and predicted values remained below 0.35 even at the maximum sample size (20,000 individuals). Even with this large sample size, not all cognitive phenotypes could be predicted with equal accuracy (Figure 2). Fluid intelligence was predicted with the highest correlation coefficient of up to 0.14, followed by processing speed and numeric memory with ∼0.1 when using *LCOR* after removing age, gender, and TIV. The prediction accuracy of processing speed was higher than that of the other three cognitive phenotypes when using age, gender, and TIV only (Scenario 4, see Figure 1). However, as shown in the black colored curves of Figure 2, the predictability of fluid intelligence, visual memory, and numerical memory scores was close to each other.

### Age, gender, and TIV result in higher accuracy than rsfMRI features

Next, we tested how age, gender, and TIV predict cognitive performance when used as sole input features and without any rsfMRI data involved (Figure 1-D.1). As shown in Figure 2 and Supplementary Figure S1, this scenario resulted in the highest correlation between actual and predicted targets across all sample sizes, outperforming all scenarios where rsfMRI features were utilized (Figure 1-D.1 to D.3). When individual characteristics served as input features, the sample size required to reach the plateau was also substantially lower (less than 500 subjects; see Figure 2). In other words, the ability of individual characteristics to predict cognitive phenotypes from a small sample size was better than the ability of rsfMRI features to predict the same targets, even when a larger sample size was used.

Given that the individual characteristics outperformed rsfMRI features in predicting cognitive phenotypes, the next logical step was to combine each TC and FC features with individual characteristics and see if it improves the predictions. For all rsfMRI features, this scenario produced the highest prediction accuracy of the first three analysis scenarios (Figure 2C, see also Supplementary Figure S1). The distinction between combined rsfMRI features and individual characteristics (Figure 2, Scenario 3) and rsfMRI features only (scenarios 1 and 2) was more pronounced when predicting processing speed in comparison to the other three cognitive phenotypes.

### tSNR plays no major role

We then asked whether excluding brain regions with high noise levels would increase prediction accuracy. We used a group-level tSNR map to threshold the rsfMRI feature maps (see Methods). Figure 3 shows fluid intelligence prediction accuracies for scenarios 1 to 3 after stepwise thresholding on the tSNR maps from 0% (no threshold, Figure 2) to 60% with 5% increments. Prediction accuracies improved with increasing sample size and the number of suprathreshold ROIs. The finding was consistent across the other cognitive phenotypes (see Supplementary Figures S3–S5). Prediction accuracy for fish consumption remained at chance-level for all tSNR thresholds (Supplementary Figure S6).

**Figure 3:**
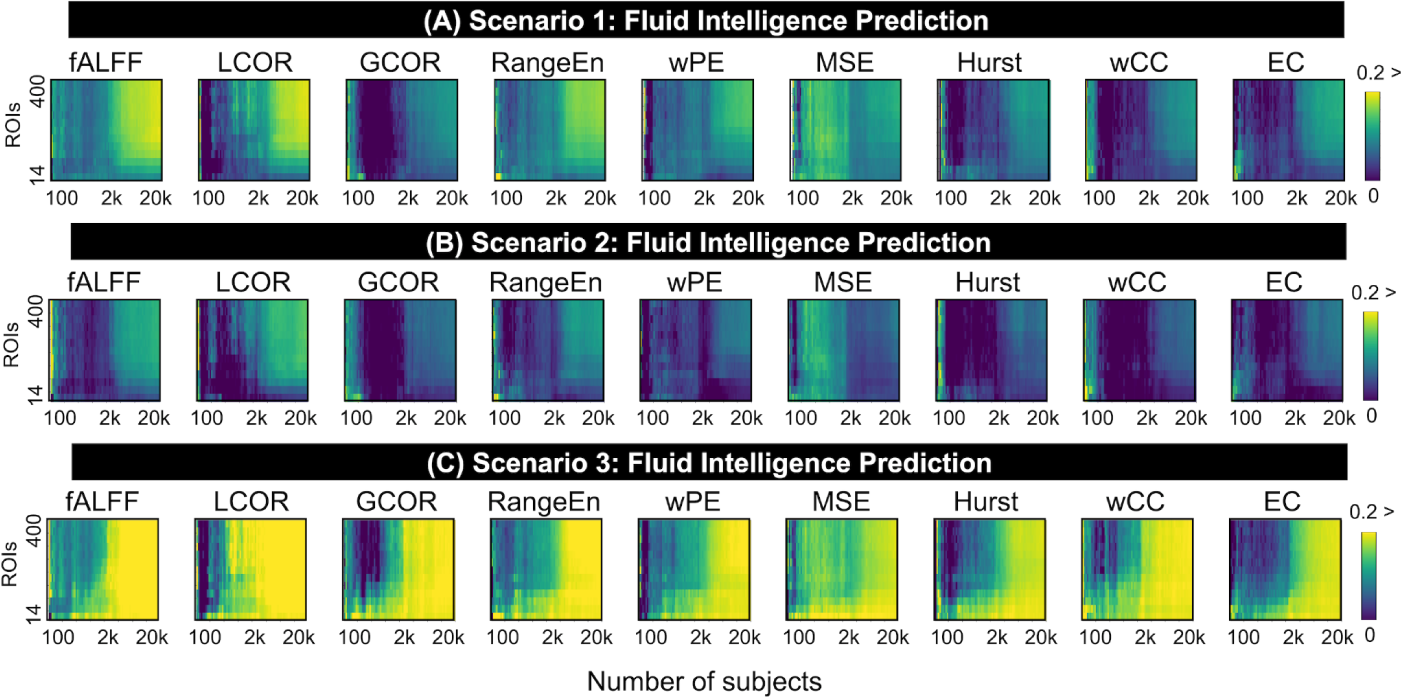
Pearson correlation accuracy associated with ridge regression modeling of *fluid intelligence* using nine rsfMRI features and after tSNR thresholding from 0% (no threshold) to 65%. In each figure panel, the accuracy values are color-coded. Additionally, the x-axis represents the population size in the analysis, and the y-axis shows the number of suprathreshold ROIs after tSNR thresholding. The predictive modeling of each pair of features and targets is repeated for different sample sizes in the UK Biobank, ranging from *N_subject_* = 100 to *N_subject_* = 20,000. The population sizes from 100 to 2000 were increased with a 50-step increment, and from 2000 to 20,000 with a 500-step increment.

### Age and gender are easier to predict than cognitive phenotypes

We investigated the capability of the rsfMRI features to predict age and gender. *wPE* and *RangeEn_B_* (TC) performed best at large sample sizes, as well as *fALFF* and *LCOR* (FC), with correlation coefficients of up to 0.5. This accuracy was considerably better than the prediction accuracy of cognitive phenotypes, which was typically less than 0.25 (see Figure 2 versus Figure 4). This result was noticeably different when individual characteristics were used as features for predictive modeling (gender and TIV for age prediction, and age and TIV for gender prediction). Gender could be classified using age and TIV with 88% accuracy. However, gender and TIV did not perform well in age prediction (ρ = 0. 2).

**Figure 4:**
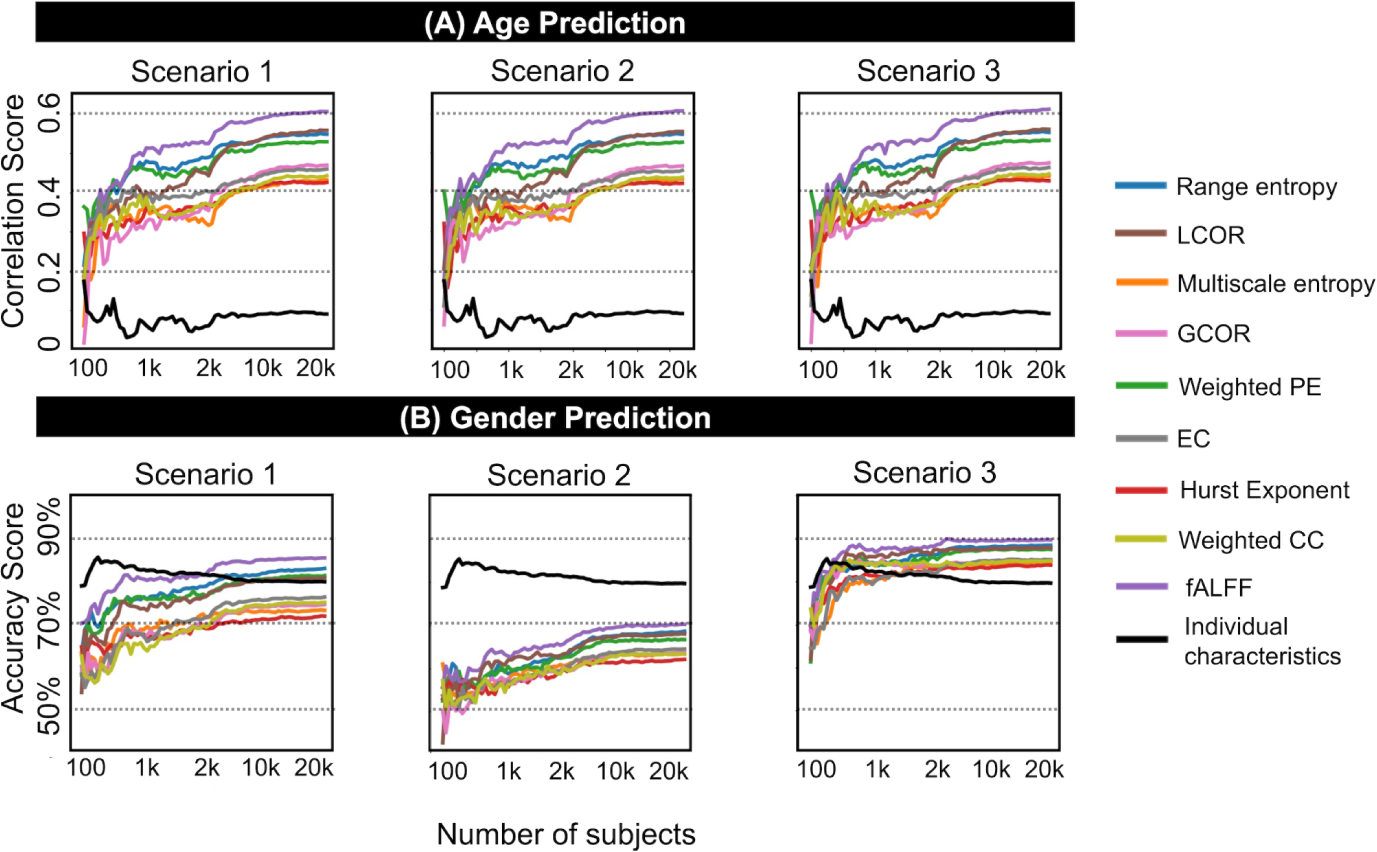
Prediction accuracy scores associated with nine rsfMRI features and age and gender as targets using scenarios 1–3 of this study (see also Figures 1-B.1–B.3 and Methods). The prediction accuracies of individual characteristics only (Scenario 4 in Figure 1-B.4) have been plotted in bold black on all panels. For age prediction, we considered gender and TIV as confounds, while for gender prediction, we considered age and TIV as confounds. Age prediction accuracies are computed as the Pearson correlation between the actual values and predicted values through ridge regression modeling. Gender prediction accuracies are computed as the balanced accuracy through ridge binary classification. Each rsfMRI feature is illustrated in a distinct color and listed in the figure legend. In each figure panel, the x-axis represents the population size in the analysis, and the y-axis shows the prediction accuracy. The predictive modeling of each pair of features and targets is repeated for different sample sizes in the UK Biobank, ranging from *N_subject_* = 100 to *N_subject_* = 20,000. The population sizes from 100 to 2000 were increased with a 50-step increment (see the light orange shadow in the figure panels) and from 2000 to 20,000 with a 500-step increment (see the light blue shadow in the figure panels). See Supplementary Figure S2 for the boxplot representation of these results.

### Similar individual patterns across rsfMRI features

We observed that some rsfMRI features have comparable predictive capacity, despite their mathematical definitions and interpretations being quite different. For instance, *fALFF* and *wPE* were frequently among the most predictive features across all analysis scenarios. Therefore, we quantified the similarity between rsfMRI features using an individual *identification* paradigm (see Methods). A number of rsfMRI feature pairs showed a high level of match across subjects (Figure 5). The pairs *wCC*-*EC*, *wPE*-*RangeEn_B_*, *fALFF*-*LCOR*, and *MSE*-*HE* were among the most highly matched. The identification accuracy increased when age, gender, and TIV were either removed or when rsfMRI features were combined with individual characteristics (Figure 5, panels B, C, and D). Importantly, identification accuracy decreased as the number of subjects increased, as opposed to the increase in prediction accuracy (Figure 2).

**Figure 5:**
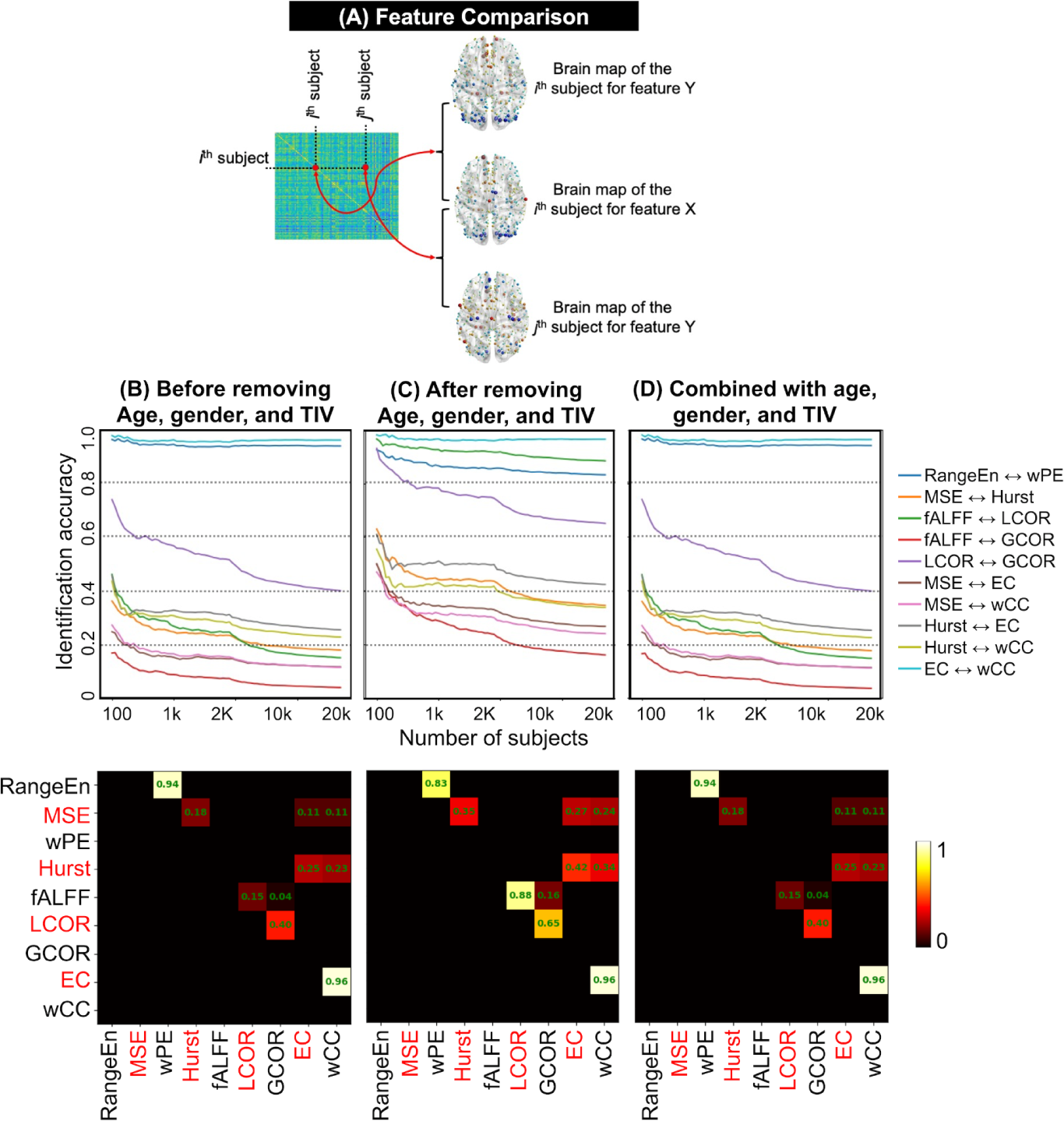
The process and results of rsfMRI feature comparison. (A) A schematic example of comparing two rsfMRI features *X* and *Y* from the same subject in a sample. This comparison leads to the computation of an identification accuracy score (see Methods). (B-D) Identification accuracy patterns of 10 rsfMRI feature pairs with above zero matching are associated with three analysis scenarios of this study (see Figure 1 as well as Methods). Each pair in the middle row panels has been depicted in a distinct color, and all pairs are listed in the figure legend. In each figure panel, the x-axis represents the population size in the analysis, and the y-axis shows the identification accuracy. The identification analyses are repeated for different sample sizes in the UK Biobank, ranging from *N_subject_* = 100 to *N_subject_* = 20,000. The population sizes from 100 to 2000 were increased with a 50-step increment (see the light orange shadow in the figure panels) and from 2000 to 20,000 with a 500-step increment (see the light blue shadow in the figure panels). The color-coded matrices in the row illustrate the identification accuracy of rsfMRI feature pairs for *N_subject_* = 20,000.

## Discussion

A primary goal of neuroscience is to investigate the relationship between brain dynamics and individual differences in behavior [56]. Spontaneous fluctuations in blood oxygenation level-dependent (BOLD) changes measured by fMRI have been shown to exhibit complex dynamics in the time domain referred to as TC [57], [58]. The interactions between BOLD changes across brain areas, also known as FC, provide useful perspectives on brain activity at a large scale. It has been demonstrated that these functional interactions are crucial for accomplishing tasks and are related to cognitive phenotypes [59].

The goal of the present study was to investigate the capacity of TC and FC features to predict cognitive phenotypes and to learn more about the effects of individual characteristics in this procedure. To this end, we looked at how well four cognitive phenotypes, fluid intelligence, processing speed, and visual/numeric memory characteristics, can be predicted by various aspects of TC/FC. We first demonstrated that, despite having different mathematical definitions, the TC and FC features lead to comparable performance across a wide range of sample sizes [28].

Comparing MRI modalities for cognitive prediction has been the subject of several recent studies [60]–[62]. However, many of these studies have utilized the same datasets, primarily the widely used Human Connectome Project (HCP) database [63], with a medium sample size of fewer than 1500 participants. Cognitive prediction studies that use the UK Biobank database have a lot more participants than HCP, but the number of studies using large fMRI datasets is still limited in the literature [22], [54], [55]. This makes extrapolating the findings of small sample size studies to larger sample sizes challenging [5]. Additionally, repeated usage of the same database in several studies (such as HCP) may result in the problem of *dataset decay*, which means that repeated statistical tests on the same dataset in different studies may result in a rise in false positives [64]. Studies on reproducible brain-wide associations have also been established to require the involvement of thousands of participants [23]. To take these issues into account, we used a sizable portion of the UK Biobank database [28], varied the sample size from 100 to 20,000 subjects, and examined the effects of sample size scaling on the predictive capacity of rsfMRI TC and FC-derived measures for cognitive phenotypic prediction. Altering the sample size indeed showed a significant impact on the predictive accuracy (Figures 2 and 4). This finding implies that prediction accuracy derived from small populations may not be reliable and can lead to large variance in cross-validation [65]. Previous cognition prediction studies using the UK Biobank and other rsfMRI databases also found that the accuracy of ridge regression modeling reaches a plateau with increasing sample size [54], [66]. The comparative predictive ability of the TC and FC features of rsfMRI (Figures 2 and 3) suggests that the complex dynamics of rsfMRI and spontaneous functional interaction between brain regions carry complementary information about behavior and cognition rather than being two distinct facets of brain function [59]. Such a relationship has also been seen in the characteristics of neurological conditions such as epilepsy, which has been described as both a disorder of functional networks in the brain and an abnormality of its dynamics at the same time [67].

Depending on the analysis workflows of this study, we either used rsfMRI features, individual characteristics, or both as input features for cognitive prediction. Interestingly, the prediction accuracy using age, gender, and TIV was higher than that of all rsfMRI features (Figure 2). This result is consistent with a previous study in the ADHD-200 Global Competition, which found that when performing classification of ADHD diagnostics, individual characteristic data (site of data collection, age, gender, handedness, performance IQ, verbal IQ, and full scale IQ) performed better than a variety of fMRI features [26]. Overall, the three individual characteristics performed better than the combined feature vectors of size 403 and the 400 rsfMRI feature vectors. A possible reason for this is that the number of samples needed to estimate a model with a given level of accuracy rises exponentially with the number of features, i.e. *the curse of dimensionality* [68]. However, given our large sample size, this is unlikely to be a concern.

Previous studies have shown that some cognitive phenotypes can be predicted better than others using neuroimaging data [5]. This is supported by our prediction results (Figure 2 and Supplementary Figure S1), which show that regardless of the rsfMRI features used and the sample size, the processing speed measure was usually predicted better than the visual and numerical memory scores. A recent review of human fluid intelligence prediction using neuroimaging data has reported an average correlation of 0.15 with a *CI*_95%_ of [0.13, 0.17] across the fMRI literature [9]. This is confirmed by our fluid intelligence prediction results with a maximum correlation score of 0.23 using combined *LCOR* and individual characteristics and at very high sample sizes (20,000 - see Figure 2). Contrary to cognitive phenotypes, age and gender were easier to predict using both TC and FC features, as shown by a comparison between the prediction accuracy curves (Figures 2 and 4 as well as the boxplots in Supplementary Figures S1 and S2). However, gender prediction using age and TIV was better than age prediction using gender and TIV (see Figure 4). All four analysis workflows passed a sanity check using the chance-level prediction of yesterday’s fish consumption (Figure 2 and Supplementary Figure S1).

The issue of noise and artifacts can influence fMRI features, for example in *fALFF* which utilizes bandpass filtering of fMRI time series [44]. tSNR is a metric for comparing the strength of an interest signal to the amount of background noise in the time domain [69]. The tSNR analysis results of our study (Figure 4, Supplementary Figures S3-S6) indicate that even the rsfMRI features of brain regions with a relatively low tSNR, which are typically found in deeper areas of the brain and close to sinus cavities, still contain predictive information about cognition. It is corroborated by our observation that using more brain regions, even when their tSNR is rather low, leads to higher accuracy. To retain the same number of brain regions across individuals, we used the group-mean tSNR map of the full sample with 20,000 UK Biobank individuals to threshold subject-specific rsfMRI feature brain maps. This is because tSNR brain maps do not always agree on the same brain regions across participants. We believe that the information in this group-mean tSNR map from a very large sample is so compressed and dimensionally reduced that any influence of data leakage would be minimal.

Our results show a relatively inverse association between identification accuracy and prediction accuracy across different sample sizes. While adding more subjects improved cognitive phenotypic prediction accuracy (Figure 2), doing so reduced identification accuracy (Figure 5). This shows that the identification problem becomes harder as the sample size grows because there are more chances of obtaining a match with another subject than with the self. On the other hand, the prediction problem becomes relatively easier for larger populations because more information is available for learning. Additionally, it is consistent with a recent study that found a similar dichotomy between the neural identity facets that best predict a person’s behavior and cognition and those that best distinguish them from other people [61]. Our results also suggest that the nine investigated rsfMRI features can be categorized into various matched pairs (Figure 5). High identification accuracy between *wPE* and *RangeEn_B_*, *HE* and *MSE*, *fALFF* and *LCOR*, and *wCC* and *EC*, are most notable. The similar prediction performance of these feature pairs can be partly explained by the measurements yielding similar individual-level patterns (as shown by the high identification accuracy), even though they are conceptually different.

In conclusion, rsfMRI TC and FC measures show potential for cognitive, phenotypic, and individual characteristic prediction. However, age, gender, and TIV showed a higher capacity for predicting cognitive phenotypes than these rsfMRI features. Having said that, there are several limitations that should be taken into account before arriving at a solid conclusion. First, we only used one predictive modeling algorithm ridge regression, so other models might capture different information. Second, even though many more variables, such as handedness and genetic factors, could influence the rsfMRI features, we only considered three individual characteristics in our predictive modeling. Third, we attempted to address the challenge of accurately quantifying cognitive phenotypes using some of the most reliable cognitive phenotypes available in the UK Biobank [53]. Despite this, these quantitative scores might still be unreliable and subject to oversimplification [70]. Unfortunately, standardized normed scores that account for demographic factors such as age and gender are not available in the UK Biobank database. Fourth, we only included cortical areas in our analyses. Using subcortical and cerebellar areas may provide a more complete picture. Taken together, our findings could aid future research in creating more accurate, individualized models of cognitive prediction.

## Methods

### Data and preprocessing

We used the rsfMRI data of 20,000 unrelated UK Biobank (UKB) participants after excluding subjects with mental and cognitive disorders (ICD10, category F), diseases of the nervous system (ICD10, category G), and cerebrovascular diseases (ICD10, categories I60 to 69). Data management of the UKB datasets was performed using DataLad [71] on JURECA, a pre-exascale modular supercomputer operated by the Jülich Supercomputing Center at the Forschungszentrum Jülich, Germany. The duration of each rsfMRI scan was 6 minutes (490 time points), with a repetition time (*TR*) of 0.735 sec, an echo time (*TE*) of 39 msec, a voxel size of 2.4×2.4×2.4 mm, and a field of view of 88×88×64. The following procedure was performed on the rsfMRI datasets as part of a pipeline developed on behalf of the UK Biobank [52]: grand-mean intensity normalization of the entire 4D fMRI dataset by a single multiplicative factor; highpass temporal filtering using Gaussian-weighted least-squares straight line fitting with *σ* = 50 sec; echo planar imaging unwarping; gradient distortion correction unwarping; and structured artifact removal through independent component analysis (ICA), followed by an ICA-based X-noiseifier (ICA-FIX) [72]–[74]. No spatial or temporal smoothing was applied to the fMRI volumes. The preprocessed data files, referred to as “*filtered_func_data_clean.nii”* in the UK Biobank database, were normalized to the MNI space using FSL’s *applywarp* function with spline interpolation and parcellated using the Schaefer brain atlas into 400 ROIs [75]. Since we needed a continuous fMRI time series for the extraction of TC features, we did not apply motion scrubbing. Finally, we considered age, gender, and TIV as individual characteristics in the analyses and incorporated them into four analysis scenarios illustrated in Figure 1. The TIV of each subject was extracted after brain extraction from the T1 image using the Computational Anatomy Toolbox (CAT12) for SPM [76].

Four cognitive phenotypes were selected as the predictive targets among the most reliable UK Biobank cognitive phenotypes, including fluid intelligence (data field 20016), processing speed (data field 20023), numeric memory (data field 20240), and visual memory (data field 399) [53]. Additionally, an unrelated binary target (fish consumption yesterday - data field 103140) was used as a sanity check of the rsfMRI features in the predictive modeling scenarios.

### Temporal complexity features

*HE* [48] is used to determine whether a time series contains a *long-memory process*. It quantifies three different types of trends: (*i*) values between 0.5 and 1, indicating that the time series is complex and has long-range dependence; (*ii*) values less than 0.5, indicating that the time series is random and has short-range dependence; or (*iii*) a value close to 0.5, indicating that the time series is a random walk with no memory of the past. HE has been shown to be stable and reproducible across different fMRI datasets [77]. In this study, we estimated *HE* using the rescaled range analysis technique [48]. The *wPE* [49] is a modified version of permutation entropy [78], that captures order relations between time points in a signal and generates an ordinal pattern probability distribution using an embedding dimension *m* and a time delay *τ,* where the former is the length of the patterns and the latter is a lag parameter denoting the number of time points to shift throughout the time series. In this study, we used the parameters *m* = 4 and *τ* = 1 and normalized the *wPE* values by dividing them by *log*_2_(*m*!) in order to get the numbers between 0 and 1. *RangeEn* offers two versions (*RangeEn_A_* and *RangeEn_B_*) as modifications to approximative entropy [79] and sample entropy [80], respectively. A property of *RangeEn_B_* is that regardless of the nature of the signal dynamics, it always reaches 0 at its tolerance value of *r* = 1 [50]. In light of this, one can obtain a complete trajectory of signal dynamics in the *r*-domain using this measure. Therefore, we extracted this trajectory from ROI-wise rsfMRI and reduced its dimensionality by computing the area under each curve along the *r*-axis (*m* = 2). We have already shown that range entropy is robust to variations in signal length [50], making it a viable option for relatively short-length time series such as rsfMRI. *MSE* is an extension of sample entropy that provides insights into the complexity of rsfMRI fluctuations over a range of time scales [51]. The measure returns a trajectory of sample entropy values across the time scales 1 to *τ_max_*. We have shown that *MSE* of rsfMRI may be linked to higher-order cognition [37]. In this study, we chose the parameters *m* = 2, *r* = 0.5, and *τ_max_* = 10 for *MSE*. We then reduced its dimensionality by taking the area under their curves and dividing by *τ_max_*.

### Functional connectivity features

We computed the FC measures of rsfMRI at two spatial scales: (*i*) at the ROI level (*EC*, *wCC*) and (*ii*) first at the voxel level, then averaged within the ROIs (*fALFF*, *LCOR*, *GCOR*). For the ROI-wise measures, we characterized the FCs between every pair of ROIs in each rsfMRI dataset and extracted the connections using Pearson correlation between mean fMRI time series [47]. *GCOR* serves as a representative of brain-wide correlation properties and a voxel-level representation of node centrality [46]. *LCOR* measures voxel-level local coherence defined as the average of the correlation coefficients between a voxel and its immediate surroundings (a Gaussian kernel with FWHM of 25 mm) [45]. Similar to *GCOR*, *LCOR* takes both the strength and sign of functional connections into consideration. *fALFF* quantifies the contribution of low frequency fluctuations to the total frequency range within a given frequency band (here, 0.008-0.09 Hz [44]). While *GCOR and LCOR* assess the strength of interregional and local cooperation by measuring the temporal similarity between voxels, *fALFF* evaluates the amplitude of regional neuronal activity. For each subject, the voxelwise *GCOR*, *LCOR*, and *fALFF* brain maps were parcellated into 400 ROIs using a brain atlas [75]. *EC* is an ROI-based measure that indicates the impact of an ROI on the functional brain network [47]. The EC of the *i*^th^ ROI corresponds to the *i*^th^ element in the eigenvector corresponding to the largest eigenvalue of the ROI-wise functional connectome. *wCC* quantifies how much the ROIs in the brain network functionally cluster together. This metric is calculated as the ratio of all triangles in which the *i*^th^ ROI participates to all triangles that, theoretically, could be formed given the degree of the *i*^th^ ROI’s involvement in the brain’s functional network [47]. The list of rsfMRI features in this study is summarized in Table S1 in the Supplementary Materials.

### tSNR analysis

We calculated tSNR for each brain region as the ratio between the mean and standard deviation of its rsfMRI time series [69]. This led to a tSNR brain map for each participant, which we normalized over ROIs and later averaged across the entire UK Biobank population (*N_subject_* = 20,000). We used the group-average map for thresholding to exclude the *noisiest* brain regions at multiple tSNR levels. The thresholding levels were varied from no threshold (i.e., preserving all ROIs for the prediction) to 65%, resulting in 14 suprathreshold ROIs. See Figure 3 as well as Figures S1 to S4.

### Predictive modeling

Following previous studies in the field [5], [54], [55], we chose to use ridge regression with l2-norm regularization and classification for predictive modeling in this study. As illustrated in Figures 1-B.1 to B.4, we designed four analysis scenarios based on the role of individual characteristics in predictive modeling. We trained 78 ridge regression models for each cognitive phenotype on a wide range of UK Biobank subjects from *N_subject_* = 100 to *N_subject_* = 2000 with a 50-step increment and from *N_subject_* = 2000 to *N_subject_* = 20,000 with a 500-step increment and each tSNR level, resulting in a total number of 36504 models (9 features × 4 targets × 78 population sizes × 13 tSNR levels). We also trained 78 ridge binary classifiers using each rsfMRI feature to predict fish consumption yesterday (total number of models: 9126). In all cases, we estimated the best model hyperparameter *λ* of the ridge regression/classification over the following values: [0.001, 0.01, 0.1, 1, 5, 10, 100, 1000, 10000, 100000] through grid search. For the evaluation of prediction accuracy, we performed five repeats of 5-fold nested cross-validation using the scikit-learn [81] and Julearn (https://juaml.github.io/julearn/main/index.html) libraries in Python. For evaluation of the regression models, we computed Spearman correlation coefficient between the actual targets and the model’s predictions. To evaluate the binary classifications, we used balanced accuracy, taking into account any imbalance between the two classes. We repeated the predictive modeling for five targets (four cognitive phenotypes as well as fish consumption yesterday) and nine rsfMRI features at a range of sample sizes varying from 100 to 20,000. At each sample size, we randomly sampled the data to contain an equal number of males and females. We developed four predictive modeling scenarios based on the role of age, gender, and TIV in our study, as illustrated in Figure 1. These scenarios included (*B.1*) prediction using rsfMRI features before removing individual characteristics, (*B.2*) prediction using rsfMRI features after treating individual characteristics as confounds and removing them, (*B.3*) prediction using combined rsfMRI features and individual characteristics, and (*B.4*) prediction using individual characteristics only. Individual characteristics were regressed out at the target level for regression modeling [5], [7] and at the feature level for the classification analyses [24], [82] using linear regression. Confound removal was performed in a cross-validation consistent manner to avoid data leakage [24].

### Feature comparison via identification analysis

We adapted the individual *identification paradigm* from the functional connectome fingerprinting literature [10], [83] and applied it to comparing different rsfMRI features of the same subject across a population. In this context, “identification” refers to the process of identifying a rsfMRI feature vector (brain map) *X* having the highest spatial correlation with *Y,* one of the other eight rsfMRI feature maps across the entire population. The identification accuracy was defined as the proportion of correctly identified individuals based on matching their two rsfMRI features. The score ranges between 0 and 1, with higher values indicating a better match. See Figure 5-A for a schematic example of comparing two rsfMRI features across a given sample.

## Acknowledgments

This work was supported by the Schwerpunktprogramm (SPP2041), project number 454012190, EI 816 28-1 (Machine-learning on Brain Connectomics: Individual Prediction of Cognitive Functioning in Health and Cerebral Small Vessel Disease), and the Helmholtz Portfolio Theme “Supercomputing and Modelling for the Human Brain”. SBE acknowledges funding by the European Union’s Horizon 2020 Research and Innovation Program (grant agreements 945539 (HBP SGA3) and 826421 (VBC)) and the Deutsche Forschungsgemeinschaft (DFG, SFB 1451 & IRTG 2150). All data analyses for this study were performed on the JURECA pre-exascale modular supercomputer operated by the Jülich supercomputing center at Forschungszentrum Jülich, Germany.

## Author contributions

Each author made a substantial contribution to this study. The final draft of the manuscript was read and approved by all authors. We use the CRediT contributor role taxonomy to describe individual contributions to the paper. Conceptualization: A.O., L.S., D.I.L., S.B.E., K.R.P.; Data Curation: A.O., L.S., F.H., J.K., J.D.; Formal analysis: A.O.; Funding acquisition: S.B.E., K.R.P., G.T., B.C.; Methodology: A.O., L.S., S.B.E, K.R.P; Software: A.O., L.S., F.R., K.R.P.; Supervision: S.B.E., K.R.P.; Visualization: A.O.; Writing—original draft: A.O.; Writing—review & editing: A.O., L.S., D.I.L., F.R., F.H., J.K., J.D., M.P., G.T., B.C, S.B.E., K.R.P.

## Competing interests

GT has received fees as consultant or lecturer from Acandis, Alexion, Amarin, Bayer, Boehringer Ingelheim, BristolMyersSquibb/Pfizer, Daichi Sankyo, Portola, and Stryker outside the submitted work. The remaining authors declare no conflicts of interest.

## Data availability

UK Biobank data can be obtained via its standardized data access procedure (https://www.ukbiobank.ac.uk/).

## Supplementary Materials

**Table S1.** RsfMRI features and the prediction targets in this study

| RsfMRI Feature name | Category | Spatial resolution |
| --- | --- | --- |
| Weighted permutation entropy (*wPE*) [49] | Temporal complexity | ROIwise |
| Range entropy (*RangeEn<sub>B</sub>*) [50] | Temporal complexity | ROIwise |
| Multiscale entropy (*MSE*) [51] | Temporal complexity | ROIwise |
| Hurst exponent (*HE*) [48] | Temporal complexity | ROIwise |
| Eigenvector centrality (*EC*) [47] | Functional connectivity | ROIwise |
| Weighted clustering coefficient (*wCC*) [47] | Functional connectivity | ROIwise |
| Fractional amplitude of low-frequency fluctuations (*fALFF*) [44] | Functional connectivity | Voxelwise |
| Local correlation (*LCOR*) [45] | Functional connectivity | Voxelwise |
| Global correlation (*GCOR*) [46] | Functional connectivity | Voxelwise |
| Prediction target | UK Biobank Data field |  |
| Fluid intelligence | 20016 |  |
| Processing time | 20023 |  |
| Visual memory | 399 |  |
| Numeric memory | 20240 |  |
| Fish consumer yesterday | 103140 |  |

**Figure S1:**
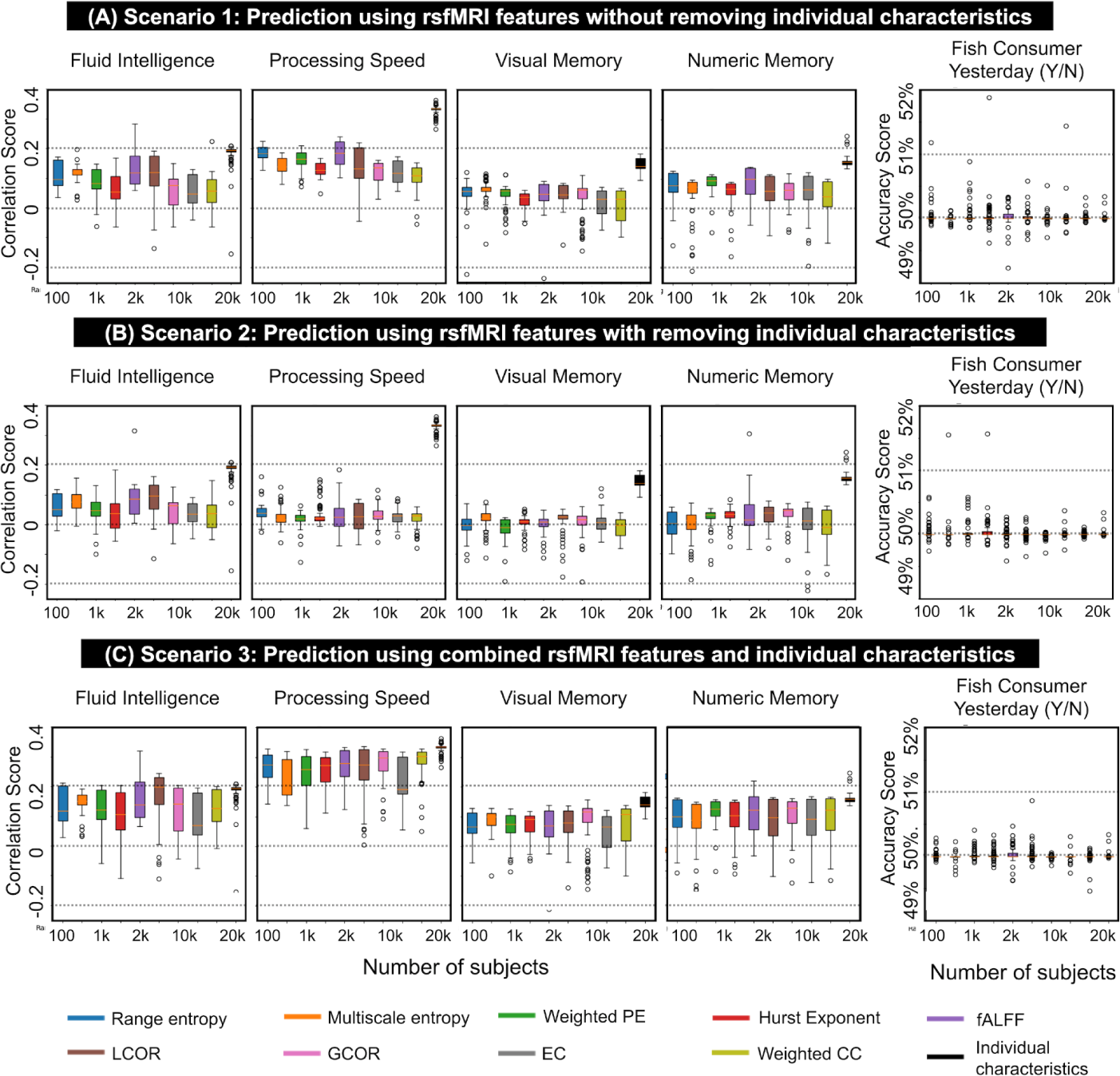
Prediction accuracy scores associated with nine rsfMRI features and five prediction targets using scenarios 1–3 of this study (see also Figures 1-B.1–B.3 and Methods). Prediction accuracies of the fluid intelligence, processing speed, visual memory, and numeric memory scores are computed as the Pearson correlation between the actual values and predicted values through ridge regression modeling. The prediction accuracy of Fish consumer yesterday is computed as the balanced accuracy through ridge binary classification. Each rsfMRI feature is illustrated in a distinct color and listed in the figure legend. In each figure panel, the box has a line at the median and spans the complete range of sample sizes (from 100 to 20,000 participants), extending from the lower to upper quartile values of the prediction accuracies. The whiskers extend outside the box to display the data’s range. The population sizes from 100 to 2000 were increased with a 50-step increment and from 2000 to 20,000 with a 500-step increment. See Figure 2 for the representation of prediction accuracies over the range of sample sizes.

**Figure S2:**
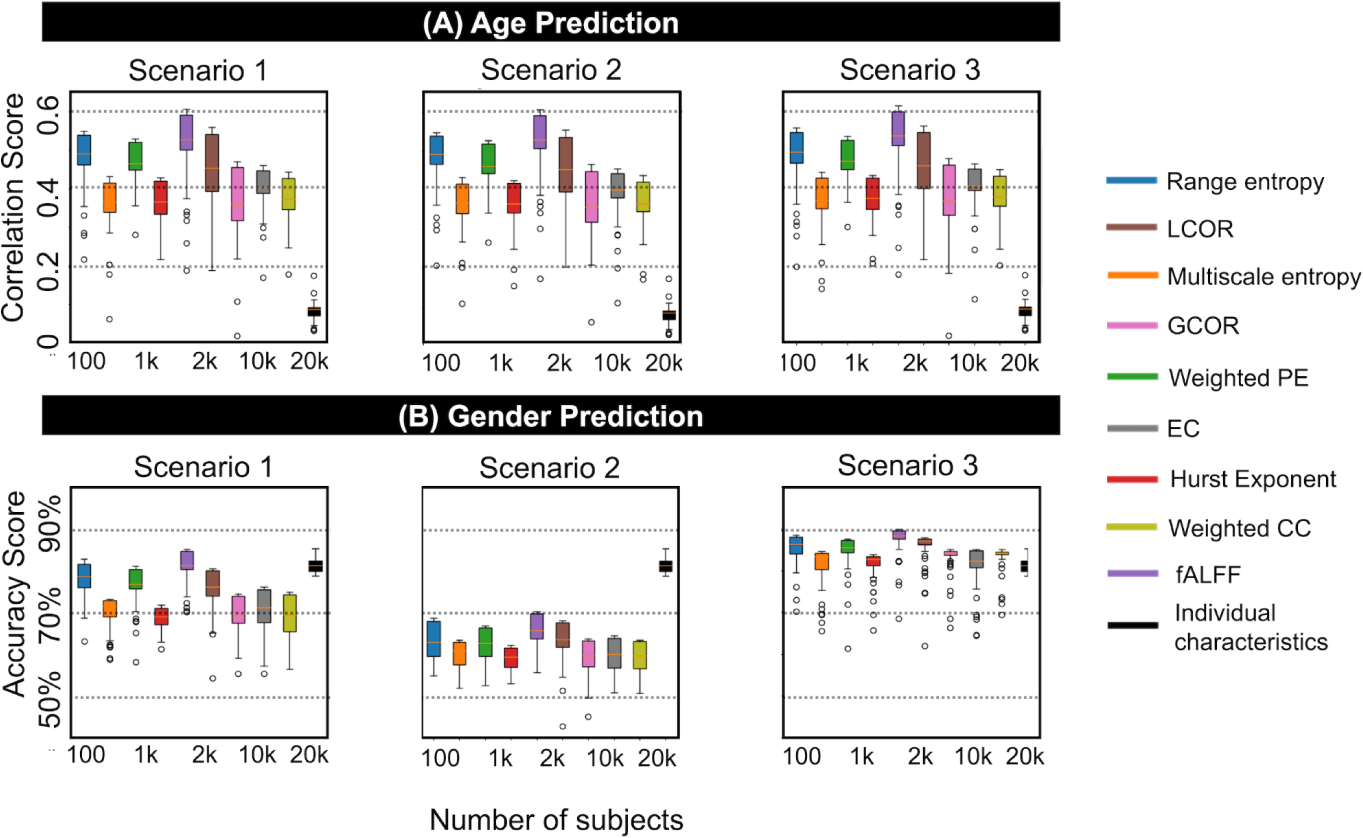
Prediction accuracy scores associated with nine rsfMRI features and age and gender as targets using scenarios 1–3 of this study (see also Figures 1-B.1–B.3 and Methods). For age prediction, we considered gender and TIV as confounds, while for gender prediction, we considered age and TIV as confounds. Age prediction accuracies are computed as the Pearson correlation between the actual values and predicted values through ridge regression modeling. Gender prediction accuracies are computed as the balanced accuracy through ridge binary classification. Each rsfMRI feature is illustrated in a distinct color and listed in the figure legend. In each figure panel, the box has a line at the median and spans the complete range of sample sizes (from 100 to 20,000 participants), extending from the lower to upper quartile values of the prediction accuracies. The whiskers extend outside the box to display the data’s range. The population sizes from 100 to 2000 were increased with a 50-step increment and from 2000 to 20,000 with a 500-step increment. See Figure 4 for the representation of prediction accuracies over the range of sample sizes.

**Figure S3:**
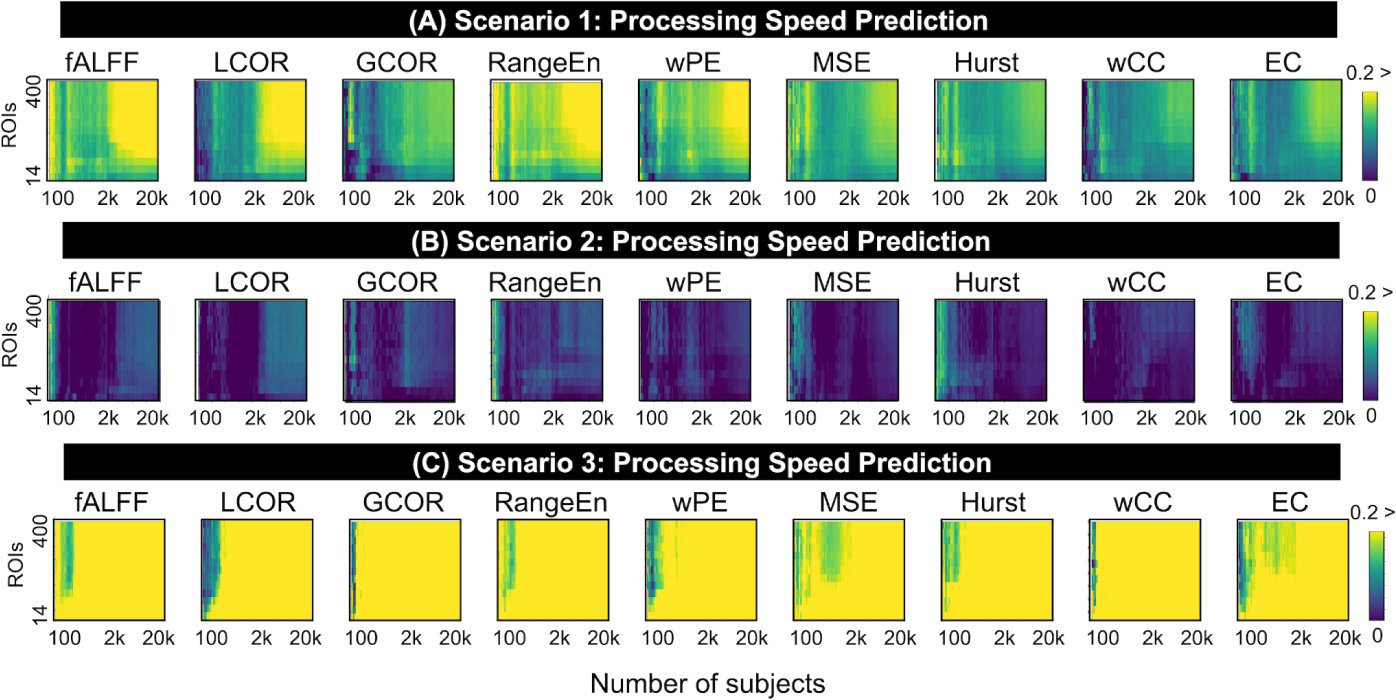
Pearson correlations associated with ridge regression modeling of the *processing speed* score using nine rsfMRI features and after tSNR thresholding from 0% (no threshold) to 65%. In each figure panel, the accuracy values are color-coded. Additionally, the x-axis represents the population size in the analysis, and the y-axis shows the number of suprathreshold ROIs after tSNR thresholding. The predictive modeling of each pair of features and targets is repeated for different sample sizes in the UK Biobank, ranging from *N_subject_* = 100 to *N_subject_* = 20,000. The population sizes from 100 to 2000 were increased with a 50-step increment, and from 2000 to 20,000 with a 500-step increment.

**Figure S4:**
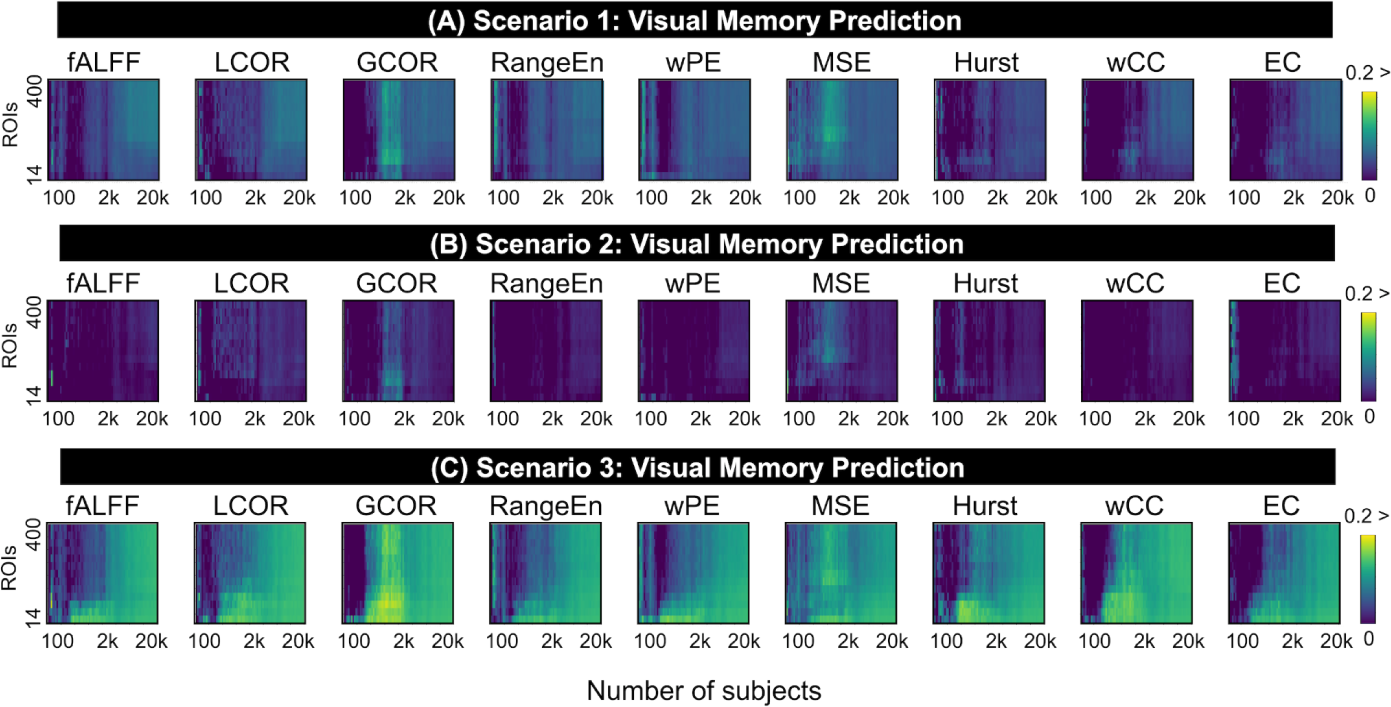
Pearson correlation accuracies associated with ridge regression modeling of the *visual memory* score using nine rsfMRI features and after tSNR thresholding from 0% (no threshold) to 65%. In each figure panel, the accuracy values are color-coded. Additionally, the x-axis represents the population size in the analysis, and the y-axis shows the number of suprathreshold ROIs after tSNR thresholding. The predictive modeling of each pair of features and targets is repeated for different sample sizes in the UK Biobank ranging from *N_subject_*= 100 to *N_subject_* = 20,000. The population sizes from 100 to 2000 were increased with a 50-step increment, and from 2000 to 20,000 with a 500-step increment.

**Figure S5:**
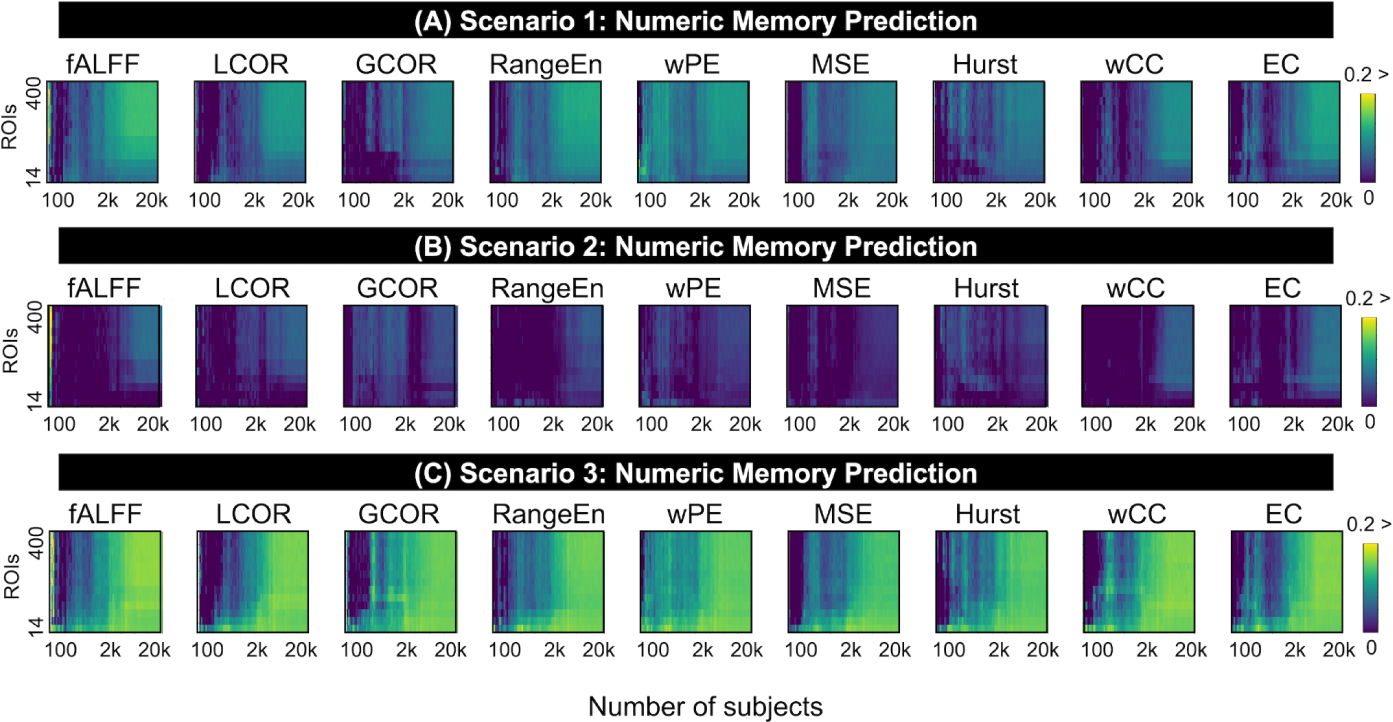
Pearson correlation accuracies associated with ridge regression modeling of the *numeric memory* score using nine rsfMRI features and after tSNR thresholding from 0% (no threshold) to 65%. In each figure panel, the accuracy values are color-coded. Additionally, the x-axis represents the population size in the analysis, and the y-axis shows the number of suprathreshold ROIs after tSNR thresholding. The predictive modeling of each pair of features and targets is repeated for different sample sizes in the UK Biobank, ranging from *N_subject_* = 100 to *N_subject_* = 20,000. The population sizes from 100 to 2000 were increased with a 50-step increment, and from 2000 to 20,000 with a 500-step increment.

**Figure S6:**
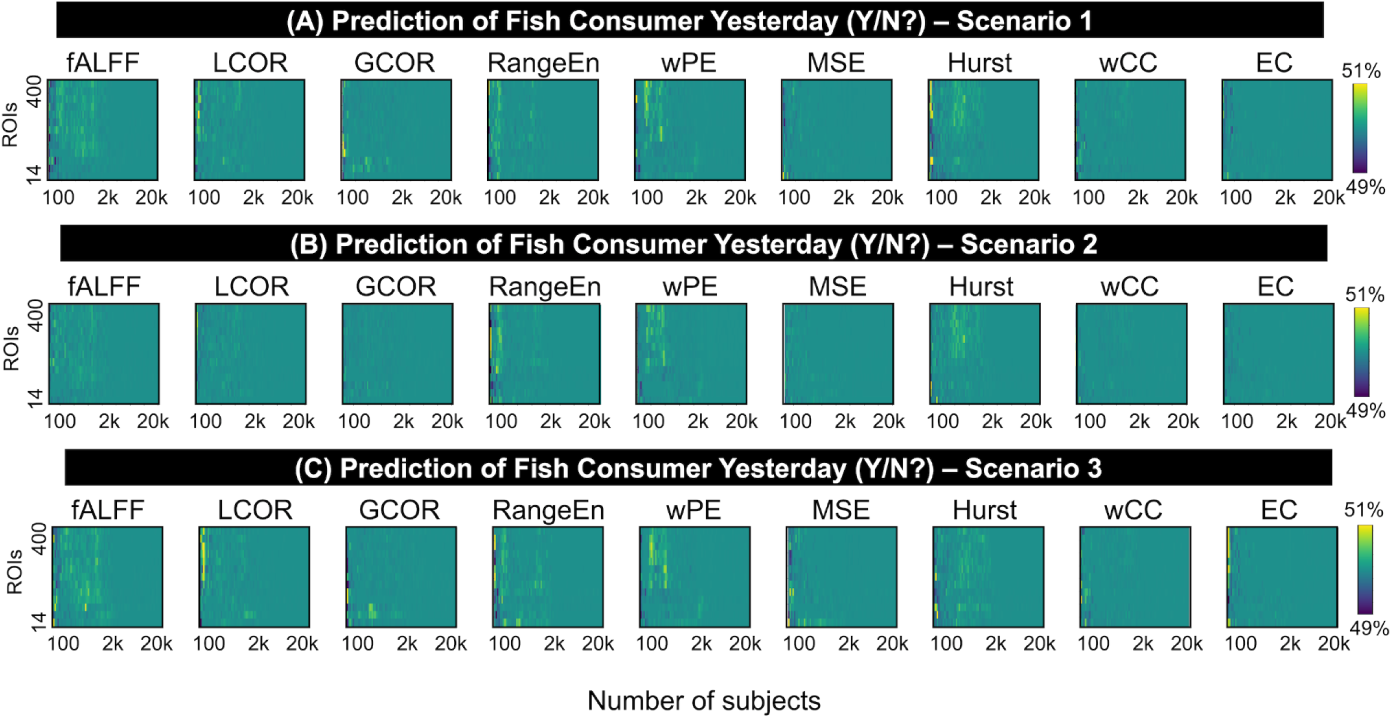
Balanced accuracy associated with ridge binary classification of *fish consumption yesterday* using nine rsfMRI features and after tSNR thresholding from 0% (no threshold) to 65%. In each figure panel, the accuracy values are color-coded. Additionally, the x-axis represents the population size in the analysis, and the y-axis shows the number of suprathreshold ROIs after tSNR thresholding. The predictive modeling of each pair of features and targets is repeated for different sample sizes in the UK Biobank, ranging from *N_subject_* = 100 to *N_subject_* = 20,000. The population sizes from 100 to 2000 were increased with a 50-step increment, and from 2000 to 20,000 with a 500-step increment.

## Notes

https://biobank.ctsu.ox.ac.uk/crystal/browse.cgi

### Summary of Updates

Editing throughout the manuscript

